# Effects of commercial unflavored and vanilla-flavored e-liquids on nicotine intake and withdrawal

**DOI:** 10.64898/2026.01.12.697777

**Authors:** Deniz Bagdas, Nardos Kebede, Jennifer Sedaille, Tore Eid, Hanno C. Erythropel, Julie B. Zimmerman, Nii A. Addy

## Abstract

Electronic cigarette liquids (e-liquids) often contain flavors and solvents that may influence nicotine addiction. In this study, we characterized the dose-response relationship of commercial unflavored nicotine e-liquids and investigated the impact of vanilla-flavored e-liquids on nicotine vapor self-administration (VSA) and withdrawal in rats. Male adolescent Sprague Dawley rats self-administered aerosols generated from commercial e-liquids containing 0, 3, 6, or 12 mg/ml nicotine in a propylene glycol (PG) and glycerol (G) vehicle. The vehicle (0 mg/ml nicotine) supported robust VSA, indicating the reinforcing effects of PG/G vapor. 3 mg/ml nicotine did not support VSA, while both 6 and 12 mg/ml nicotine concentrations produced significant reinforcement, with 6 mg/ml yielding the most stable responding. The 6 mg/ml concentration was selected for subsequent comparisons with vanilla-flavored e-liquids. Vanilla flavor (0 mg/ml nicotine) led to maintained VSA behavior, confirming its reinforcing effects. However, the combination of vanilla and nicotine (6 mg/ml) did not alter nicotine intake or withdrawal severity, as assessed by mecamylamine-precipitated somatic signs. Blood nicotine and cotinine levels were similar between nicotine and vanilla + nicotine conditions, indicating that vanilla flavor did not affect systemic nicotine metabolism. Additionally, the PG/G vehicle induced significant somatic signs, suggesting that vapor exposure itself, independent of nicotine, contributes to these physiological responses. These findings provide critical insights into the reinforcing and physiological effects of both nicotine and non-nicotine constituents in e-cigarette aerosols, underscoring the need for future studies and regulatory strategies that consider the abuse liability of flavors and solvents, such as PG/G, particularly among adolescents.

**Significance Statement:** Flavored electronic nicotine delivery systems raise concern for promoting nicotine use in youth. Using a rat vapor self-administration model, we show nicotine produces concentration-dependent reinforcement, while vanilla flavor is reinforcing but does not enhance nicotine intake or withdrawal.

## 1. Introduction

The introduction and availability of electronic nicotine delivery systems (ENDS) has significantly changed the way people consume nicotine worldwide. Examples of ENDS include vapes, vaporizers, vape pens, hookah pens, electronic cigarettes (e-cigarettes), e-cigars, and e-pipes. In particular, the use of e-cigarettes has increased dramatically over the past decade, with adolescents and young adults representing a significant proportion of users. ^1,2^ Surveys indicate that the prevalence of vaping is particularly high among adolescents, many of whom initiate e-cigarette use at an early age. ^3^ The widespread availability of flavored e-liquids, including vanilla and other sweet flavors, has been identified as a major factor driving e-cigarette experimentation and sustained use in youth. ^2,4^ Despite increasing concerns about the addiction potential of flavors in ENDS, prior investigations of the mechanisms underlying the behavioral effects of flavored nicotine exposure were limited to menthol and green apple. ^5–7^ Moreover, there is only little understanding as to whether early adolescent exposure to nicotine-containing ENDS aerosol increases susceptibility for continued ENDS use in adulthood. This gap in understanding highlights the need for further investigation.

While human studies provide valuable epidemiological insights, investigating how adolescent nicotine exposure through ENDS use increases susceptibility to continued use in adulthood, and the potential role of flavors, cannot be ethically conducted in human subjects. This highlights the urgent need for preclinical models of vapor exposure, such as vapor self-administration (VSA), to investigate the behaviors and mechanisms underlying continuous vaping behavior. Rodent models provide a controlled and mechanistic approach to study nicotine reinforcement, dependence, and the influence of flavors on self-administration. The establishment of preclinical models can also be used to examine the complex interactions of nicotine type (free base vs. salt formulations that contain a pH modifier such as benzoic acid), nicotine source (tobacco-extracted vs. synthetic), flavors, the ratio of the solvent propylene glycol (PG) and glycerol (G), and e-cigarette characteristics such as power output and temperature. Rodent models also make it possible to test commercial tobacco products in controlled laboratory conditions, providing critical insight into how product formulations and device characteristics influence nicotine use behaviors.

Vanillin flavor has been found in e-liquids labeled as vanilla-flavored and other flavored e-liquids, such as caramel or those with a label that suggests a baked good. ^8,9^ In addition, several studies have identified vanillin as one of the most prevalent flavorants in e-cigarettes. ^10,11^ Vanilla flavor is also common in, smokeless tobacco, little cigars, and cigarillos. ^8,9,12,13^ Vanilla is extracted from the *Vanilla planifolia* flower and is commonly associated with confectionery and dessert-related aromas. Its complex chemical composition includes numerous compounds, with vanillin (4-hydroxy-3-methoxybenzaldehyde) and ethylvanillin (4-hydroxy-3-ethoxybenzaldehyde) being the primary contributors to its characteristic scent. Studies have shown that e-cigarettes with vanilla flavor enhance the perception of sweetness and increase user-reported pleasantness. ^14,15^ Our recent studies in an operant oral self-administration paradigm revealed that vanilla flavor has its own reinforcing potential and enhances oral nicotine self-administration. ^16^ Because there is limited preclinical research on the impact of vanilla on nicotine use in the context of vapor exposure, investigating this relationship could provide valuable insights into its effects on nicotine VSA. Here, we not only characterized dose response curves of commercial nicotine e-liquids but also investigated the impact of vanilla flavored e-liquid on nicotine intake and nicotine withdrawal in a rat VSA model.

## 2. Materials and methods

### 2.1. Animals

Sprague Dawley male adolescent rats (post-natal day 28, weighing 100-115 g at arrival to the lab) were purchased from Charles River Laboratories, Wilmington, MA, USA. Due to the limited availability of VSA chambers (two chambers), the study was conducted in male rats only to ensure timely completion of the experimental procedures. Rats were singly housed in a temperature- and humidity-controlled husbandry room. They were provided *ad libitum* food and water on a standard 12-h light/dark cycle (from 7am to 7pm). All experiments received approval from The Yale University Institutional Animal Care and Use Committee (IACUC). Experiments were conducted according to the National Institutes of Health (NIH) Guide for the Care and Use of Laboratory Animals.

### 2.2. Drugs and chemicals

Commercial DuraSmoke electronic cigarette liquids (e-liquids) were purchased from VapinDirect (Wauwatosa, WI) and included 2 flavors: unflavored at nicotine concentrations of 0, 3, 6, and 12 mg/ml, and vanilla at nicotine concentrations of 0 and 6 mg/ml. As per the manufacturer, the e-liquids contained tobacco-extracted nicotine and e-liquids were used as received. Nicotine concentrations used in this study were chosen based on prior experiments in a mouse model, ^7^ and a dose response curve study was performed. The non-flavored 0 mg/ml nicotine e-liquid (containing only PG and G) was used as vehicle group. Nicotine content, principal flavor content, and PG/G ratio of commercial e-liquids were verified using analytical techniques (see below, and Table 1). For nicotine withdrawal experiments, mecamylamine hydrochloride (Cat no. M9020, Sigma-Aldrich, St. Louis, MO), a non-competitive nicotinic acetylcholine receptor antagonist, was dissolved in saline and administered at 1.5 mg/kg dose subcutaneously. ^17^

**Table 1.**
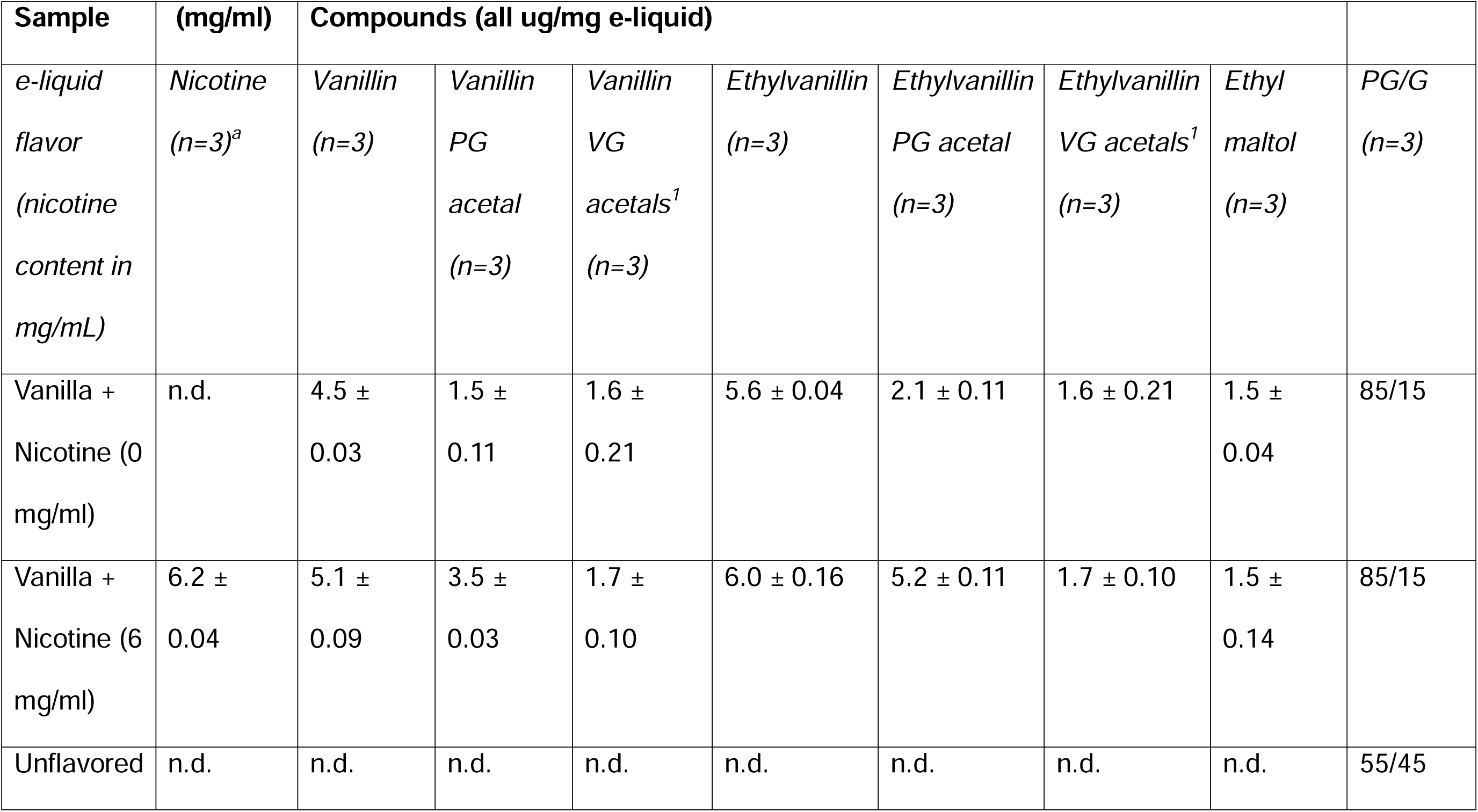
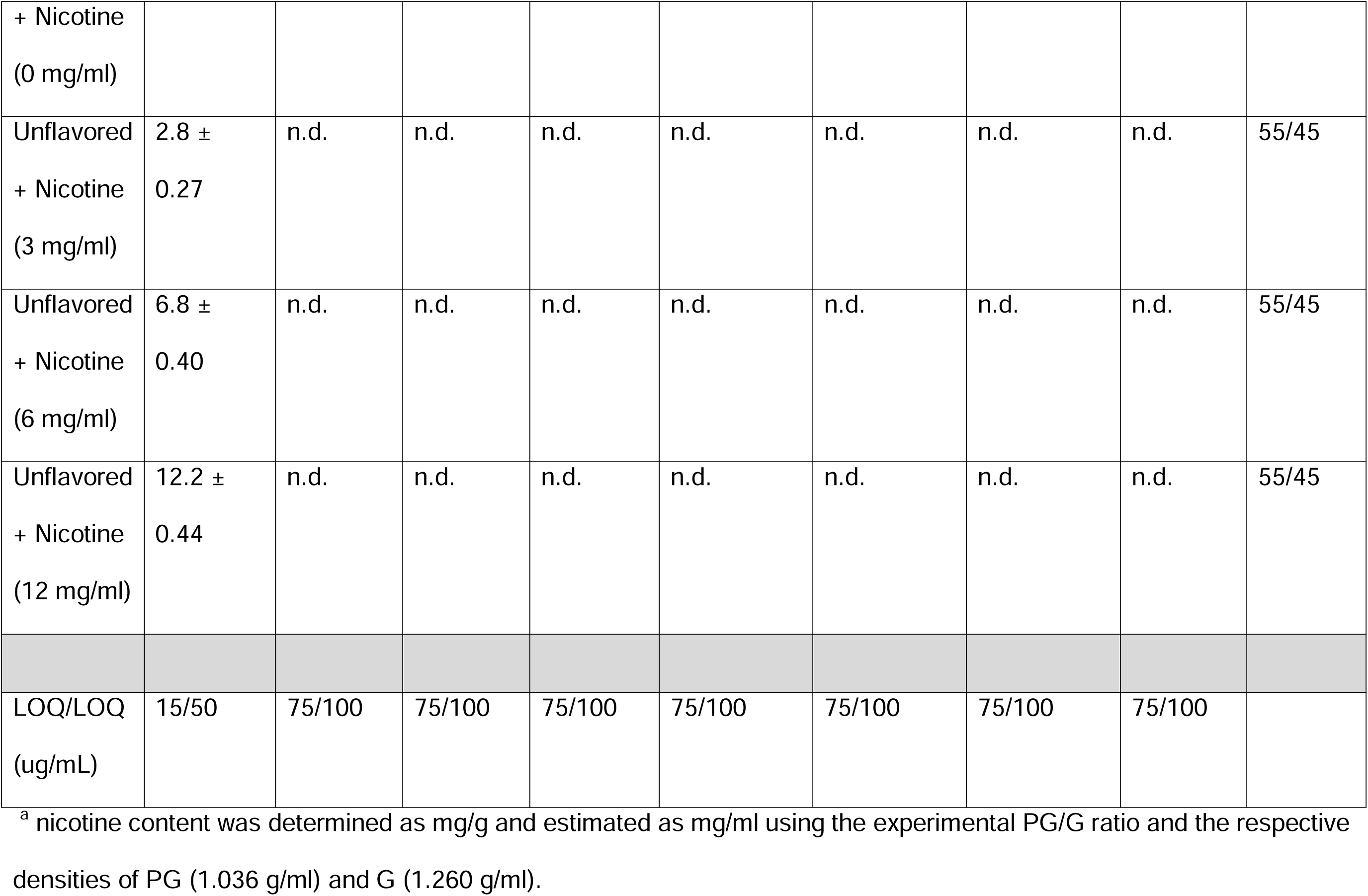
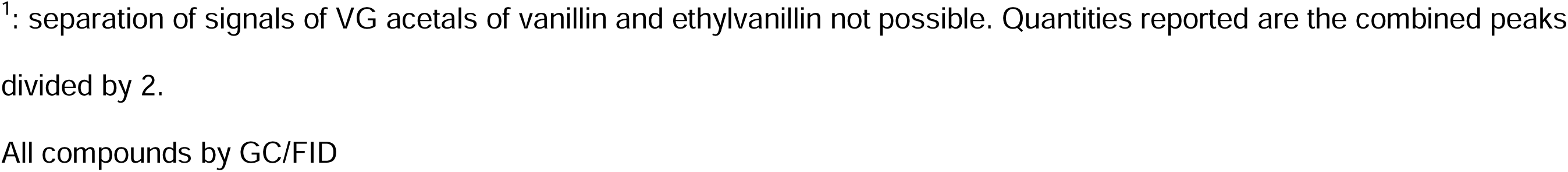
Chemical analysis for nicotine and vanilla flavor components of purchased Durasmoke e-liquids. N.d. – not detected. PG – propylene glycol. G - glycerol

Analytical standards were nicotine (99%, Alfa Aesar, Haverhill, MA), vanillin, ethylvanillin (both 99%, Sigma-Aldrich), vanillin PG acetal (mixture of isomers=97%), ethylvanillin PG acetal (mixture of isomers=98%, both Bedoukian, Danbury, CT), ethyl maltol (≥99%, Sigma-Aldrich), propylene glycol (99.5%, Fisher Scientific, Waltham, MA), glycerol (99.5%, Sigma-Aldrich). For glycerol acetals (mixture of isomers) of both vanillin and ethylvanillin, which are not available commercially, in-house synthesized calibration standards were used. ^18^

### 2.3. Vapor exposure

Rats were exposed to e-liquid aerosol generated from unflavored e-liquids containing 0, 3, 6, and 12 mg/ml nicotine, or vanilla-flavored e-liquid containing 0 or 6 mg/ml of nicotine via a combined passive and vapor self-administration (VSA) procedure. The VSA procedure was performed in an e-cigarette vapor delivery system which we modified for rats, based on previous work in a mouse model in the Henderson lab, using a similar system (T2 e-vape, La Jolla Alcohol Research, La Jolla, CA) ^7,19^. Atomizers were from GeekVape (M series dual coil, 0.40 ohms, Shenzhen IVPS Technology Co., Ltd., Shenzhen, China).

The chemical content of e-liquids is provided in Table 1 and 2. In the vapor delivery system, power was set at 50W, and heating temperature was set at 400°F. Rats arrived at the facility on a Thursday and experiments began the following Monday. For 5 days, rats received a passive 3-second vapor puff every 8 minutes over 2-hour daily sessions (15 puffs/day). This passive exposure was performed to promote the subsequent acquisition of self-administration behavior, where animals would need to reach a criterion of 2:1 active:inactive nose pokes, as previously described ^20^. The passive exposure was followed by 18 days of VSA (3-second puff, 30-second timeout, for 1 hour session daily) using fixed-ratio schedules of reinforcement (FR1 for 6 days, FR3 for 5 days, and FR5 for 7 days). Active nose poke responses were paired with vapor delivery, according to the FR schedule. Experiments were performed weekly for 5 consecutive days, from Monday to Friday, with no weekend sessions.

#### 2.3.1. Chamber habituation, side assignment, and exclusion

Vapor self-administration chambers were equipped with two nose-poke ports located on the left and right sides. The vapor delivery port could be connected to either the left or right side of the chamber, and during habituation, both the vapor port and the active nose-poke port were randomly assigned to either side, allowing for any combination of same-side or opposite-side placement. To assess potential side bias and to try and prevent reinforcing or avoidance behaviors, rats first underwent 1-2 days of habituation. During this period, they were placed in the chamber where they received non-contingent vapor puffs on one side. Nose-poke activity was recorded to identify behavioral patterns and side biases.

In some cases, rats inserted their nose into the nose-poke port on the same side of vapor delivery port and remained there for extended periods, resulting in a high number of recorded responses that did not reflect true operant behavior. To better assess side bias and engagement, rats that completed a second day of habituation were tested with the vapor port positioned on the opposite side from day one. Rats were classified as showing avoidance if more than 80% of their responses occurred at a single port, with minimal engagement (<20%) at the opposite port (high responding on one port interpreted as passive avoidance, and low responding on the other indicating a lack of true exploration or engagement). Rats showing avoidance were excluded from the study. For VSA experimental sessions (after assessment for possible side bias), all remaining rats were randomly assigned to left- or right-active nose-poke ports to ensure overall group counterbalancing. To standardize the vapor delivery port side across animals, the vapor delivery port was placed on the side opposite the assigned active nose-poke port for all rats.

### 2.4. Evaluation of nicotine withdrawal

In rodent models, mecamylamine (a non-selective nicotinic acetylcholine receptor antagonist) is commonly administered to, as it precipitates withdrawal symptoms in nicotine-exposed animals by blocking nicotinic receptors and unmasking physical signs of dependence.

Following nicotine VSA, rats were evaluated for mecamylamine-precipitated nicotine withdrawal-related behaviors. 24 hours following the last vapor exposure, rats were injected with mecamylamine (1.5 mg/kg, subcutaneously). ^17^ After 10 minutes, rats were placed in a clear acrylic cage and observed for mecamylamine-precipitated somatic signs of withdrawal (paw and body tremors, head shakes, body shakes, backing, jumps, curls, foot licks, writhes, teeth chattering, cheek tremors, yawns, and ptosis) for 20 minutes. ^21^

### 2.5. Measurements of blood nicotine and cotinine

Immediately after last vapor exposure, animals were anesthetized using isoflurane (4%) delivered via an induction chamber until anesthesia was confirmed by the absence of responses to a pain stimulus test. Blood was then collected via cardiac puncture within 5 minutes after animals were removed from VSA chambers and immediately transferred into lithium heparin-coated tubes (green-top tubes) on ice to prevent clotting. Samples were centrifuged (2000 rpm, 4°C, 15 min). Following centrifugation, 200 µl of plasma was transferred into Eppendorf microcentrifuge tubes and stored at −80°C until analysis.

Unconjugated plasma cotinine and nicotine concentrations were assayed utilizing liquid chromatography tandem mass spectrometry (LC/MS/MS, Waters Xevo TQ-XS, Waters Corporation, Milford, MA) with stable isotope (deuterated)-labeled internal standards. The assay was similar, but not identical, to that described in the literature. ^22^ Cotinine, cotinine-D3, nicotine and nicotine-D4 were purchased from Cerilliant, Round Rock, TX. All primary standards and controls were prepared in human plasma determined to be free of nicotine and cotinine. Assays were performed in a building in which tobacco use was not allowed. Separate ion transitions were used for purposes of quantitation and confirmation for each of the three analytes as follows: (1) cotinine: 177 → 80, quantifier 177 → 98; qualifier: cotinine-D3 180 → 80 internal standard; (2) nicotine: 163 → 132; quantifier, 163 → 106; qualifier, nicotine-D4 167 → 136 internal standard. The assay utilized a plasma ‘crashing’ (zinc sulfate protein precipitation/methanol extraction) technique before LC/MS/MS. Between-day coefficients of variation (CVs) over the range of values observed in this study were 6.6 to 8.1% for cotinine and 8.6 to 13.4% for nicotine.

### 2.6. Chemical analysis of commercial products

Gas chromatography (GC) was carried out for e-liquid characterization using GC-mass spectroscopy (MS), and quantification of select e-liquid components using GC-flame ionization detection (FID) using established methods ^23^. Limits of detection/quantification (LOD/LOQ) is listed in Table 1.

GC-MS was carried out on a Clarus 580 GC with SQ8S MS using an Elite-5MS column (length 60 m, id 0.25 mm, 0.25 μm film; all PerkinElmer, Waltham, MA).

Weighed samples were diluted in 1ml methanol (Thermo Scientific, Waltham, MA) and 1 μL was injected via an autosampler. The injector temperature was 300 °C with a split ratio of 10 using He as carrier gas (Airgas, Radnor, PA) and the following program was used: 40 °C for 7 min; ramp 10 °C/min to 50 °C then hold for 20 min; ramp 10 °C/min to 310 °C then hold for 8 min; ramp 10 °C/min to 325 °C then hold for 11.5 min. The MS was used in electron impact ionization (EI+) with an m/z range of 30-620.

GC-FID was carried out on a GC-2010 (Shimadzu, Columbia, MD) using a J&W DB-5 column (length 60 m, id 0.25 mm, 0.25 μm film; Agilent, Santa Clara, CA). Weighed samples were diluted in 1ml methanol (Thermo Scientific) containing 1 g/L internal standard, and 1 μL was injected via an autosampler. The injector temperature was 250 °C with a split ratio of 300 using He (Airgas) as carrier gas and the following program was used: 30 °C for 7 min; ramp 10 °C/min to 50 °C then hold for 20 min; ramp 10 °C/min to 310 °C then hold for 12 min. Detector temperature was 325 °C. Calibration curves for nicotine, vanillin, vanillin PG acetal, vanillin glycerol acetals, ethylvanillin, ethylvanillin PG acetal, ethylvanillin glycerol acetals, and ethyl maltol with ≥ 4 points were constructed for quantification. Findings were listed in Table 1.

### 2.7. Statistical analyses

We used GraphPad Prism software (version 10; GraphPad Software, Inc., San Diego, CA) to make graphs and perform statistical analyses. Prior to statistical analysis, the Bartlett’s test was used to confirm homogeneity of variances across groups. A two-way repeated measures analysis of variance (RM ANOVA with Greenhouse-Giesser correction) was used to evaluate the impact of e-liquids on the taking behavior of each test over time. We examined each e-liquid by session interaction. If the interaction was found to be significant, we performed *post-hoc* Sidak multiple comparisons among the e-liquid by session point, to determine when differences in taking behavior emerged.

A t test was used for the number of puffs earned on 18^th^ session, and a one-way ordinary ANOVA was performed for nicotine withdrawal signs, with a Bonferroni correction for multiple comparisons. Significance was determined at p < 0.05.

## 3. Results

### 3.1. Nicotine vapor self-administration across nicotine concentrations using unflavored e-liquid

Figures 1 shows total active and inactive nose pokes per hour across 18 self-administration sessions across nicotine concentrations (0, 3, 6, and 12 mg/ml) for 8 rats per group using unflavored e-liquid.

**Figure 1.**
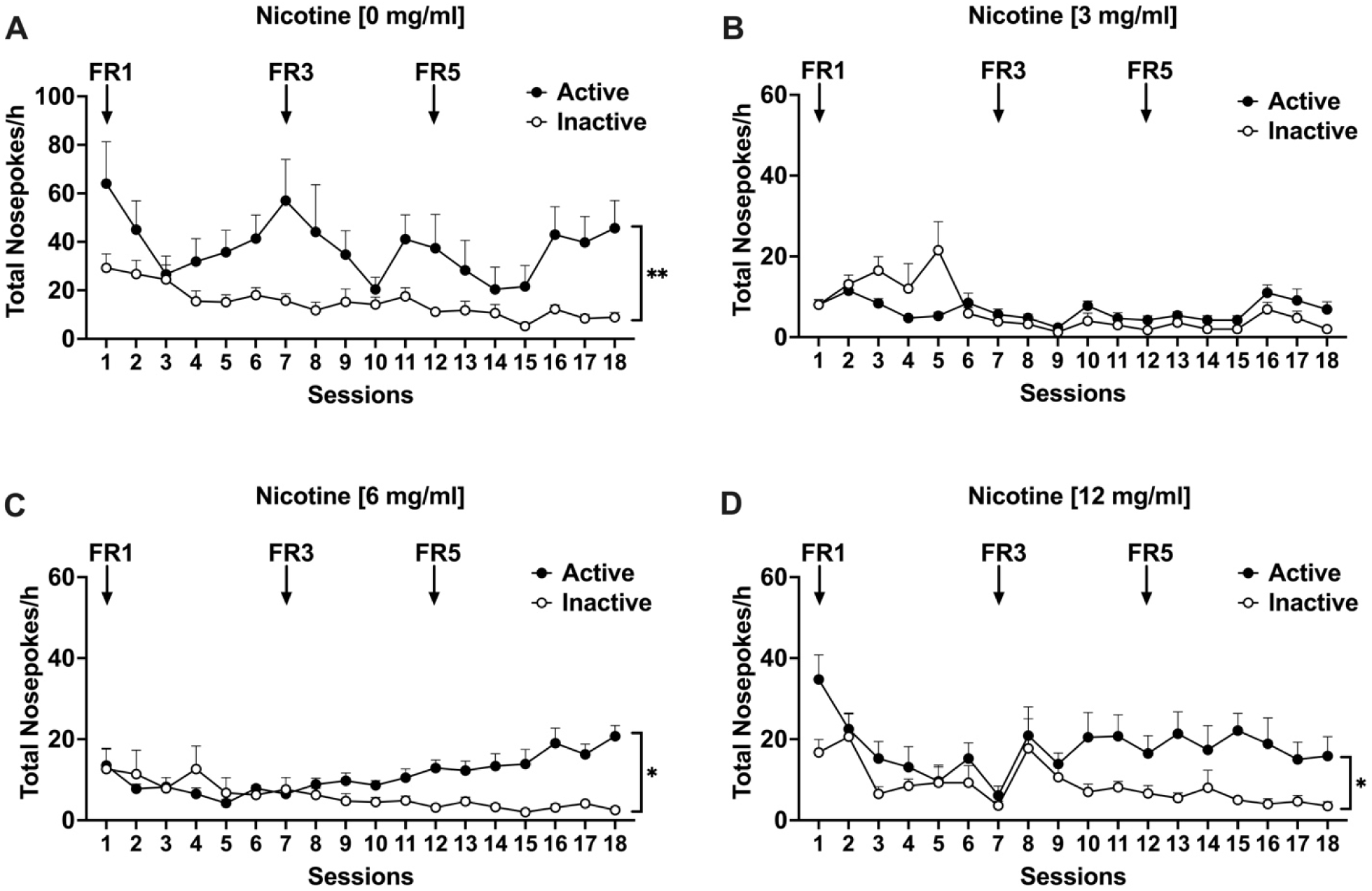
Nicotine vapor self-administration across unflavored nicotine concentrations. Total nose pokes for (A) propylene glycol and vegetable glycerin vehicle (nicotine 0 mg/ml), (B) nicotine 3 mg/ml, (C) 6 mg/ml, and (D) 12 mg/ml concentrations across 18, one-hour vapor self-administration sessions. Data expressed as mean ± standard error of the mean (SEM) of n=8/group. *p < 0.05 indicating active nose poking. FR: Fixed ratio

For the 0 mg/ml nicotine concentration (Figure 1A), active nose pokes were significantly higher than inactive responses while reaching the criteria of 2:1 active versus inactive responding [F _nose poke (1,14)_ = 9.816, p = 0.0073], but no escalation was observed across sessions [F _session (3.673,51.43)_ = 2.278, p = 0.0785; F _interaction (17,238)_ = 4.501, p = 0.4027], suggesting that while rats significantly self-administered PG/G, their response rates did not significantly change over sessions.

At the 3 mg/ml nicotine concentration (Figure 1B), there were no significant differences between active and inactive responses [F _nose poke (1,14)_ = 0.0066, p = 0.9363], and response rates remained low throughout the sessions. A significant main effect of sessions was observed [F _session (3.496, 48.94)_ = 5.399, p = 0.0017], however, the absence of significant reinforcement behavior showed that this dose was insufficient to drive vapor self-administration.

At the 6 mg/ml nicotine concentration (Figure 1C), rats reached the criteria of 2:1 active versus inactive responding. They demonstrated significantly greater responding on active nose pokes compared with inactive nose pokes [F _nose poke (1,14)_ = 8.228, p = 0.0124], without a significant effect of sessions [F _session (4.880,68.31)_ = 1.424, p = 0.2279] but with a significant nose poke x session interaction [F _interaction (17,238)_ = 4.501, p < 0.0001]. Rats showed similar active and inactive nose pokes during FR1. Beginning at FR3, however, they showed a preference for active nose pokes, and their active nose pokes increased slightly but steadily through FR3 and FR5. The results indicate that rats showed reinforced operant responding for the 6 mg/ml nicotine concentration.

In the 12 mg/ml nicotine concentration group (Figure 1D), rats showed significantly higher active nose pokes than inactive nose pokes in line with the criteria of 2:1 active versus inactive responding [F _nose poke (1,14)_ = 5.658, p = 0.0322]. Additionally, a significant main effect of sessions was observed [F _session (5.648,79.07)_ = 4.462, p = 0.0008] without an interaction with nose pokes [F _interaction (17,238)_ = 1.530, p = 0.0850]. Although no escalation was observed, early response rates declined by the end of FR1, after which rats maintained a stable self-administration pattern across FR3 and FR5 scheduling. The significant difference in responding between active and inactive nose pokes was maintained, confirming that rats showed reinforcement behavior of the 12 mg/ml nicotine concentration.

Rats self-administered nicotine vapor at different concentrations, confirmed by significant main effects of nicotine concentration [F _concentration_ (3,28) = 12.47, p < 0.0001], and a significant nicotine concentration x session interaction [F _interaction_ (51,476) = 1.499, p = 0.0176] without main effects of session [F _session_ (4.191,117.3) = 2.394, p = 0.0517; active nose pokes throughout Figure 1 A-D]. Nicotine reinforcement varied across concentrations, with higher nicotine concentrations leading to greater self-administration and the vehicle control PG/G having its own reinforcing effects.

### 3.2. Impact of vanilla flavor on nicotine vapor self-administration

Figure 2 shows the total number of active and inactive nose pokes per hour across 18 self-administration sessions in rats (n = 7/group) exposed to vanilla-flavored e-liquids, in the presence or absence of nicotine. While vanilla concentration was not labeled, it was present at both 0 and 6 mg/ml nicotine conditions. In the vanilla-only condition (Figure 2A), rats initially exhibited similar active and inactive nose pokes.

**Figure 2.**
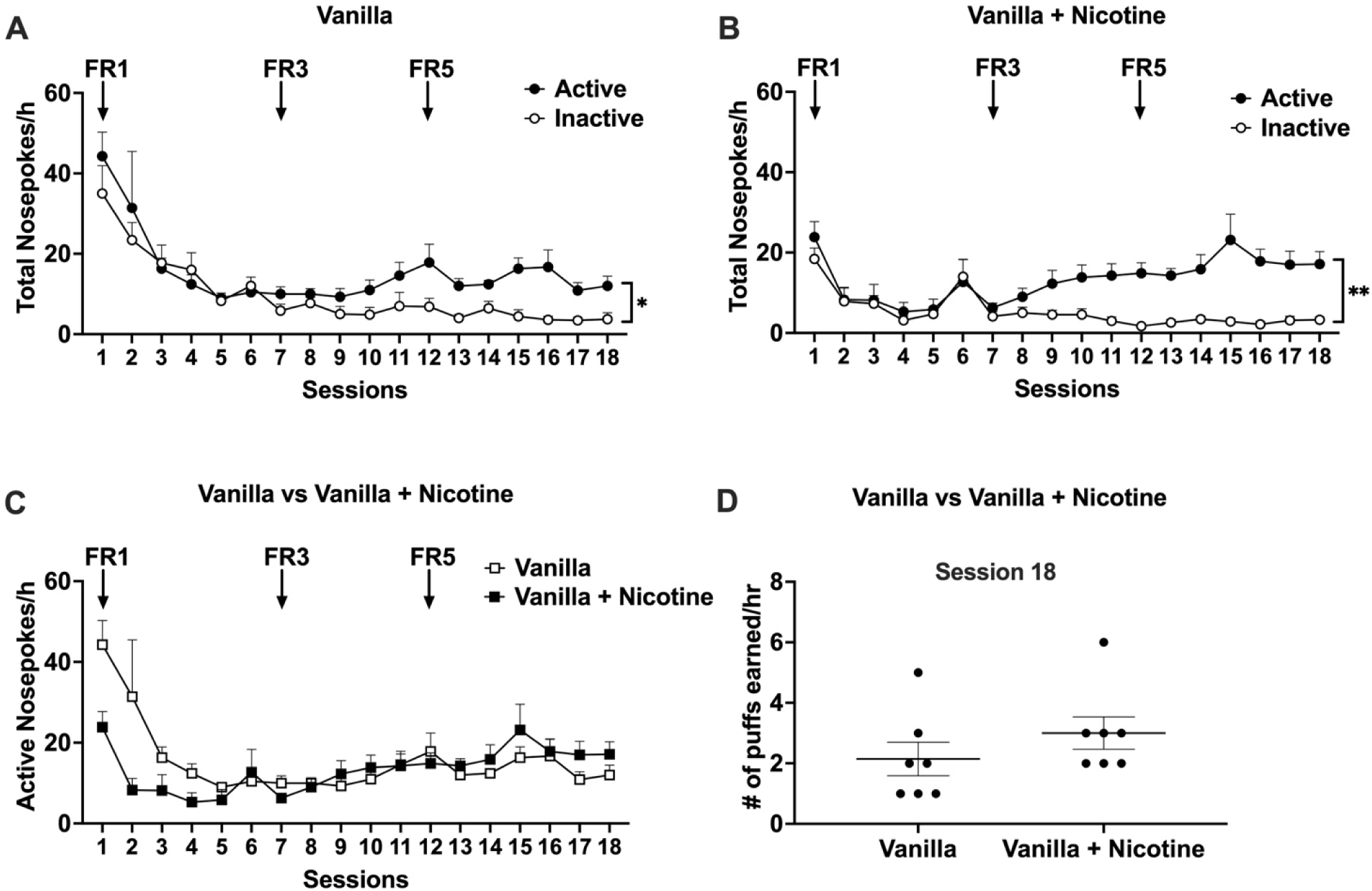
Vanilla vapor self-administration. Total nose pokes for (A) vanilla, (B) vanilla + nicotine 6 mg/ml in 18, one-hour vapor self-administration sessions. (C) Comparison of active nose poking between vanilla-alone and vanilla + nicotine groups. (D) Number of vapes per hour on 18th session of vapor self-administration. Data expressed as mean ± standard error of the mean (SEM) of n=7/group. *p < 0.05 indicating active nose poking. FR: Fixed ratio

However, as sessions progressed, active responses became significantly higher than inactive responses [F _nose poke_ (1,12) = 5.313, p = 0.0398], suggesting that the vanilla flavor alone reinforced operant responding. Both active and inactive responses were initially high during FR1 but declined over sessions. By the end of FR3, active responding became significantly higher than inactive responding, indicating a clear preference for vanilla. This pattern remained stable throughout FR5, indicating that while initial engagement was high, self-administration did not escalate but was maintained at a consistent level while reaching the criteria of 2:1 active versus inactive responding [F _session_ (2.954,35.45) = 12.46, p < 0.0001] without an interaction between nose pokes and session [F _interaction (17,204)_ = 1.030, p = 0.4271].

In the vanilla + nicotine group (Figure 2B), active nose pokes were significantly higher than inactive nose pokes [F _nose poke_ (1,12) = 15.88, p = 0.0018], confirming reinforcement and the criteria of 2:1 active versus inactive responding. A significant main effect of session [F _session_ (5.413,64.95) = 4.986, p = 0.0005], and an interaction between nose poke and session [F _interaction_ (17,204) = 3.504, p < 0.0001] were observed, with responding stabilizing across FR3 and FR5. Although no escalation in responding was observed, the difference between active and inactive responses remained significant, suggesting the reinforcing effects of vanilla-flavored nicotine.

Direct comparison of active nose pokes between vanilla and vanilla + nicotine conditions (Figure 2C) showed no significant differences in responding across sessions [F _nicotine_ (1,12) = 0.5710, p = 0.4645]. This suggests that vanilla flavor alone was sufficient to maintain vaping behavior, with no additional effect of nicotine on vapor intake. Analysis of number of vapes in the last session (Figure 2D) further supported these findings, as no significant differences in the total number of vapes were observed between the vanilla-alone and vanilla + nicotine groups (t = 1.114, df = 12, p = 0.2870). This indicates that nicotine presence did not significantly enhance vapor intake when combined with a vanilla-flavored e-liquid.

In Figure 3, we show a direct comparison of active nose pokes and vapor intake between nicotine-alone (n=8) and vanilla + nicotine (n=7) e-liquids. Active responding was compared between the nicotine-alone and vanilla + nicotine groups, revealing no significant differences in self-administration, as active nose pokes remained similar across sessions [F _vanilla_ (1,13) = 1.056, p = 0.3228; Figure 3A]. This suggests that the presence of vanilla flavor did not significantly alter nicotine self-administration behavior. Analysis of vapor intake during the final session (Figure 3B) further supports this finding, as the total number of vapes did not differ between groups (t = 1.035, df = 13, p = 0.3197). This suggests that the addition of vanilla flavoring did not further enhance nicotine intake, indicating that self-administration behavior at this nicotine concentration was maintained regardless of the presence or absence of vanilla flavoring. Moreover, blood nicotine and cotinine levels were measured in both nicotine-alone and vanilla-flavored nicotine groups (Table 2). The results showed that blood nicotine and cotinine levels were similar between groups for an average of three vapes.

**Figure 3.**
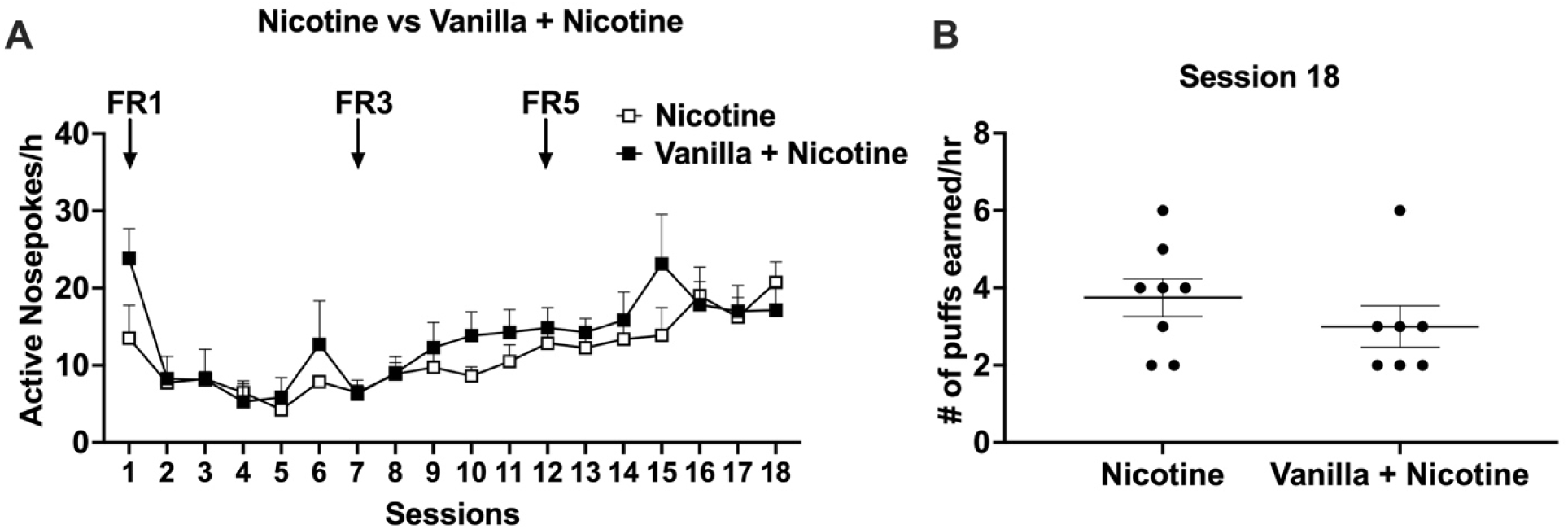
Impact of vanilla flavor on nicotine vapor self-administration. (A) Comparison of active nose poking between nicotine-alone and vanilla + nicotine groups. (B) Number of vapes per hour on 18th session of vapor self-administration. Data were expressed as mean ± standard error of the mean (SEM) of n=7-8/group. FR: Fixed ratio

**Table 2.**
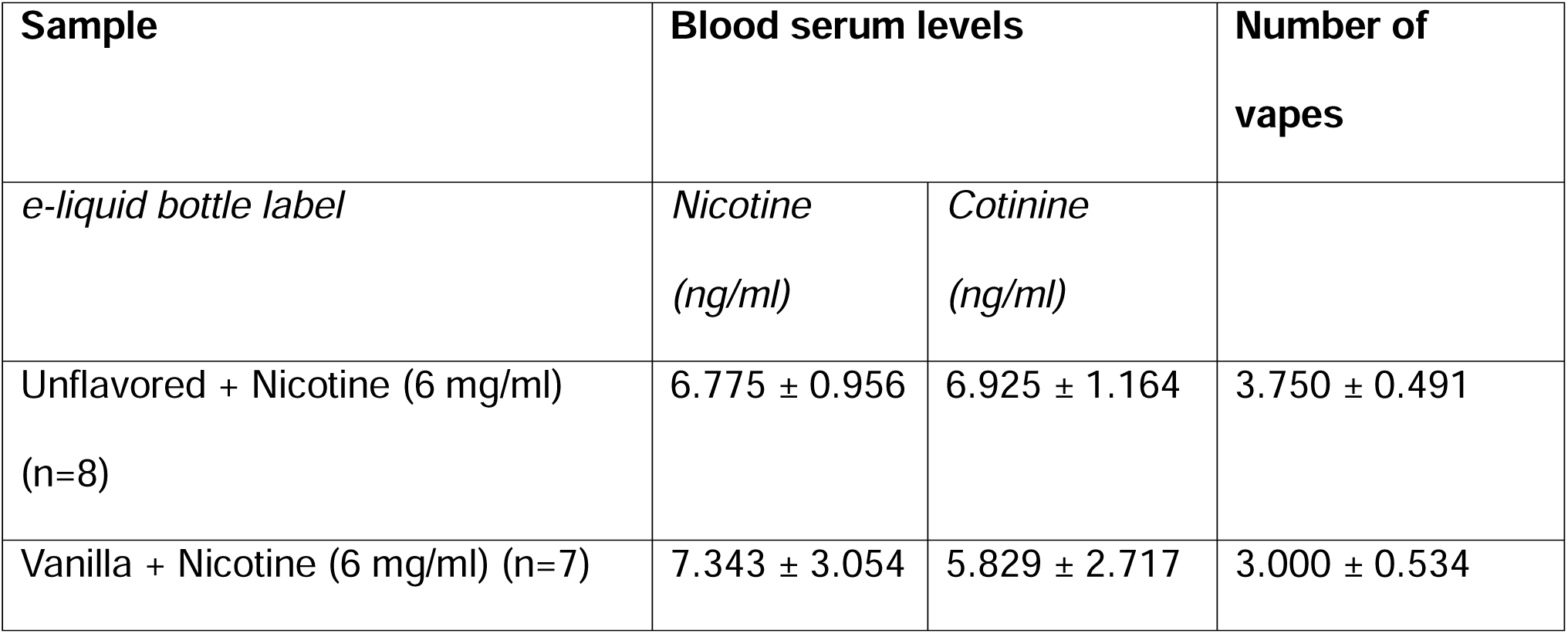
Blood nicotine and cotinine levels.

### 3.3. Mecamylamine-precipitated nicotine withdrawal in vapor self-administering rats

Mecamylamine was administered 24 hours after the last self-administration session. The number of somatic signs were quantified across experimental groups. A significant main effect of e-liquid was found [F _vanilla_ (4,33) = 20.99, p < 0.0001; Figure 4]. Naïve rats displayed minimal somatic signs. In contrast, all e-liquid aerosol-exposed groups, regardless of e-liquid nicotine content, exhibited significantly higher somatic signs compared to naïve rats (p < 0.01), indicating that vapor exposure alone contributed to these effects. Among e-liquid aerosol-exposed groups, PG/G and vanilla-alone groups showed similar levels of somatic signs (p > 0.05), suggesting that non-nicotine e-liquids were sufficient to induce these responses. Nicotine-alone and vanilla + nicotine groups displayed even higher somatic signs compared to PG/G and vanilla-alone groups (p < 0.01), confirming that nicotine further amplified these effects. However, no significant differences were observed between nicotine-alone and vanilla + nicotine groups (p > 0.05), suggesting that vanilla flavor did not alter the severity of these somatic responses.

**Figure 4.**
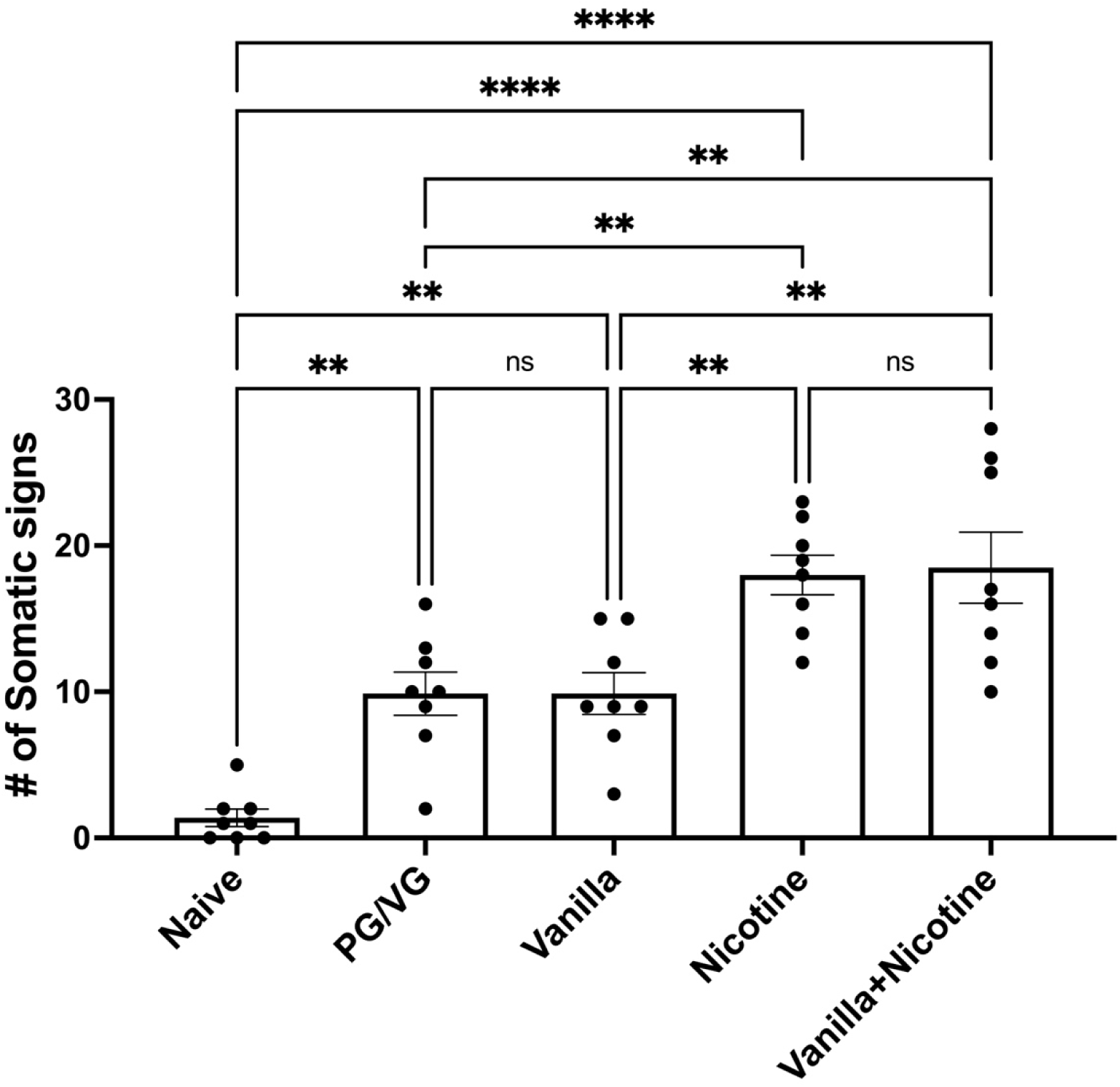
Impact of vanilla flavor on nicotine withdrawal. Mecamylamine-precipitated nicotine withdrawal following 24h vapor exposure. Somatic signs were evaluated as physical signs of withdrawal. Data expressed as mean ± standard error of the mean (SEM) of n=7-8/group. *p < 0.05 between groups.

### 3.4. Nicotine and vanilla content of the e-liquids

Nicotine and major flavorants were characterized and quantified in the tested e-liquids. The measured nicotine concentrations (2.8, 6.8, and 12.2 ug/mg e-liquid) closely matched the labeled values (3, 6, and 12, respectively), indicating accuracy in nicotine content across formulations (Table 1). Analysis of the vanilla-flavored e-liquids confirmed the presence of vanillin (4.5 – 5 ug/mg e-liquid), ethyl vanillin (5.6 – 6 ug/mg e-liquid), and ethyl maltol (1.5 ug/mg e-liquid) as major flavoring compounds (Table 1). Vanillin and ethyl vanillin are primary contributors to the characteristic vanilla aroma, while ethyl maltol provides a sweet, “cotton candy”-like note ^24^. In addition, both PG and G acetals of both vanillin and ethylvanillin were detected, which aligns with prior literature describing the reaction between these flavorants and the solvents PG and G. ^18,25,26^ PG/G ratio was found to be 55/45 for nicotine and 85/15 for vanilla e-liquids (Table 1). Unflavored e-liquids did not contain flavorants.

## 4. Discussion

In this study, we investigated the impact of aerosols from commercially available unflavored and vanilla-flavored e-liquids in the nicotine VSA. At 50W power, the unflavored 6 mg/ml nicotine concentration supported the most stable and robust VSA, indicating that this concentration was optimal for maintaining nicotine intake. Vanilla alone was also sufficient to maintain VSA behavior. The combination of vanilla and nicotine (6 mg/ml) did not significantly alter nicotine intake, suggesting that vanilla flavor did not alter the reinforcing effects of this nicotine concentration. Mecamylamine-precipitated withdrawal confirmed that nicotine exposure induced somatic signs, and that vanilla flavor did not alter nicotine withdrawal severity. Finally, blood nicotine and cotinine levels were similar between unflavored nicotine and vanilla + nicotine groups, indicating that vanilla did not alter the amount of systemic nicotine exposure. PG/G also induced VSA and significant somatic signs, suggesting that vapor exposure itself, independent of nicotine, contributes to physiological responses.

Rodent VSA models have recently gained increased attention as valuable tools for studying the behavioral effects of e-cigarette use in a controlled setting. ^7,27–29^ While the Henderson Lab employed only FR1 and FR3 schedules in mice, we modified and expanded the paradigm to FR5 to better assess nicotine maintenance in rats. For VSA, rats were tested for 18 days, beginning with 6 days on an FR1 schedule, followed by 5 days at FR3 and 7 days at FR5. This adaptation extended the protocol from the original mouse model, which utilized FR1 for 10 days, followed by FR3 for 5 days. ^7,19,20,30^ Because nicotine is a weak reinforcer in rats, the additional FR5 phase in our study allowed for further evaluation of response stability and nicotine reinforcement at higher effort requirements for rats. We also began testing with a 6 mg/ml nicotine concentration, as previous mouse models demonstrated significant nicotine VSA at this concentration. ^7^ Additionally, a study in rats reported significant VSA at 5 mg/ml nicotine, ^29^ further supporting the use of similar concentrations across species. Moreover, 6 mg/ml is a commonly commercially available nicotine concentration.

We found significant nicotine VSA (unflavored) at 6 mg/ml and subsequently tested 3 and 12 mg/ml concentrations. While 3 mg/ml nicotine did not induce VSA, both 6 and 12 mg/ml nicotine concentrations supported significant VSA behavior. Although increasing FR schedules did not lead to a robust escalation in responding, nicotine reinforcement was maintained at higher concentrations, demonstrating stable intake patterns. Our rat model effectively assesses nicotine VSA under varying concentrations and reinforcement schedules, providing a valuable tool for studying vaping behavior. Additionally, our effective concentration findings align with previous reports in male mice, ^7,19,20,30^ and male rats, ^29,31^, further supporting the reproducibility and reliability of VSA paradigms.

Rats also showed operant reinforcement behavior for the PG/G vehicle (unflavored 0 mg/ml nicotine). In rodent models, we note that prior literature findings on the reinforcing effects of PG/G alone are mixed, with some studies reporting significant reinforcement, while others suggest minimal or no reinforcing effects. ^7,29^ PG/G is often recognized as having sweet-associated taste, which may contribute to its reinforcing effects. Supporting this, clinical studies on PG and G have shown that e-cigarette users report the PG to G ratio influences palatability, flavor intensity, throat hit, and vapor density. ^32–35^

The findings with PG/G provide important insight that PG/G itself may be reinforcing and that e-cigarette characteristics such as wattage and heating temperature can influence outcomes. Indeed, a prior study revealed that PG/G alone was reinforcing when the wattage was between 40 and 55. ^28^ Given our use of 50W in the current study, our observed effects of PG/G may be attributable to this power setting. Overall, our findings are in line with prior reports that also demonstrated reinforcing effects of PG/G in mice and rats. ^28,31^ We later tested lab-made e-liquids containing PG/G alone in male adolescent and young adult rats and found similar results, but not in adult rats (data not shown). Therefore, PG/G has potential reinforcing effects in male rats at an early age at 50W power, and further research is needed to clarify the behaviors underlying its potential reinforcing effects and long-term behavioral impact.

In the current study, vanilla-flavored nicotine did not significantly alter nicotine VSA compared to nicotine alone. A recent mouse study reported that lab-made vanilla flavor alone (included 7.5 mg/ml vanillin and 7.5 mg/ml ethyl vanillin) and vanilla combined with nicotine produced similar levels of active nose pokes and vapor deliveries, and our findings are consistent with this observation. ^36^ However, we recently reported that vanillin altered oral nicotine intake, seeking behavior, and taste responses, with effects differing based on sex and vanillin concentration. ^16^ Vanillin-alone (no nicotine) induced intraoral self-administration in a concentration dependent manner in both female and male adult rats. ^16^ Vanillin also enhanced intraoral nicotine self-administration in females, increased nicotine’s hedonic taste responses, and decreased nicotine’s aversive taste responses. ^16^ We note that these findings were obtained using the two-bottle choice test, operant intraoral self-administration, and taste reactivity assays, which all assess oral nicotine consumption and palatability rather than inhaled VSA. In addition, the e-liquid used in this study contained a complex vanilla flavor formulation, not pure vanillin. This difference in composition may partly explain the discrepancy with prior oral nicotine studies where vanillin, as a single compound, has been shown to enhance intake through taste modulation. In contrast, the impact of complex vanilla flavor on VSA may be limited due to differences in sensory processing and reinforcement mechanisms between oral and inhalation routes. These findings highlight the need for further studies directly comparing the effects of isolated flavorants, such as vanillin, and complex commercial flavor formulations across different routes of administration.

This study is limited by the use of a single vanilla concentration; however, this was the only available vanilla concentration for nicotine e-liquids from this supplier. In addition, the commercial vanilla-flavored e-liquids contained several major flavor compounds besides vanillin. Moreover, the study was conducted only in male rats. The decision to use only male rats in this study was driven by logistical constraints associated with the limited availability of VSA chambers. With access to only two chambers, the study duration would have been significantly prolonged if females were incorporated. Consideration of sex as a biological variable remains important and will be addressed in future research. Lastly, vanilla flavor alone reinforced VSA behavior, and the addition of nicotine did not significantly alter response patterns. These findings highlight the potential reinforcing effects of non-nicotine e-liquid flavors and their influence on VSA behavior. Therefore, these findings also represent an important opportunity for future studies to investigate the role of PG/G and device parameters in vaping behavior.

Nicotine withdrawal is a critical component of nicotine dependence and in this study, mecamylamine-precipitated withdrawal induced significant somatic signs in nicotine self-administered rats, thereby confirming the development of physical signs of nicotine withdrawal following abstinence of VSA. Other studies have also shown the development of nicotine dependence after chronic nicotine vapor exposure as measured by nicotine withdrawal. ^17^ In adult rats, 20 mg/ml nicotine (the lowest tested concentration in the referenced study) induced both spontaneous and mecamylamine-precipitated nicotine withdrawal. ^17^ While rodents are commonly used for behavioral and mechanistic studies as a surrogate for human studies, we acknowledge that there may be behavioral and mechanistic differences in humans.

Nicotine-alone (no flavor) induced significant somatic signs compared with PG/G vehicle. Vanilla flavor (no nicotine) alone did not increase withdrawal symptoms, as somatic signs were similar between nicotine-alone and vanilla + nicotine groups, indicating no effect on nicotine dependence. In addition, after the last session of VSA, a nicotine concentration of 6 mg/ml with and without vanilla flavor induced approximately 7 ng/ml blood nicotine levels in the animals. These findings are consistent with a previous study that showed aerosol generated from e-liquid containing 3 mg/ml nicotine induced approximately 3 ng/ml blood nicotine levels in rats. ^27^ These findings suggest that while vanilla flavor can maintain VSA, it does not impact nicotine withdrawal severity. Unexpectedly, PG/G (unflavored 0 mg/ml nicotine) and vanilla-alone groups also exhibited significant somatic signs compared with naïve rats, suggesting that repeated vapor exposure, even without nicotine, may contribute to physiological responses indicative of withdrawal. Others also observed inhalation of non-nicotine containing aerosol, such as PG/G, strawberry and tobacco flavor, induced anxiety-like behaviors in light-dark transition test, as well as mecamylamine precipitation-induced decreases in locomotor activity and increases in anxiety-like behaviors in open field test. ^37^ These findings reflect an association between non-nicotine e-liquid constituent exposure and negative affect. Potential explanations for this could include sensory effects of inhalation, irritation from vapor components, or other non-nicotine factors influencing physiological adaptation. E-liquids, unlike combustible tobacco products, are available with or without nicotine. The negative physiological effects of nicotine-free aerosol inhalation are concerning, as some adolescents use non-nicotine e-liquids. ^38–40^ The potential anxiogenic effects of non-nicotine vapor inhalation require further investigation.

## 5. Conclusion

With the increasing interest in ENDS use, multiple laboratories are developing nicotine VSA models, which are a relatively new approach. These models are crucial for investigating the behavioral and physiological effects of ENDS, as they provide controlled conditions to assess nicotine reinforcement, dependence, and the impact of product constituents, such as flavors. Our findings contribute to this growing body of research, demonstrating that commercially available e-liquids with and without vanilla flavor at the same nicotine concentration can induce similar levels of self-administration. Interestingly, PG/G alone was reinforcing at 50W power, as rats self-administered it at levels comparable to, or higher than nicotine-containing e-liquids. This suggests that factors beyond nicotine, including sweet-associated sensory cues, the vehicle, or e-cigarette characteristics, such as power, may play a significant role in vapor self-administration. Further, it highlights the need for additional studies examining the role of the many non-nicotine components within e-cigarettes and their potential influence long-term product use.

## 6. Data Availability

The data underlying this study will be made available upon reasonable request to the corresponding author.

## CRediT authorship contribution statement

**Deniz Bagdas:** Writing – original draft, Supervision, Funding acquisition, Conceptualization, Methodology, Formal analysis, Data curation (Vapor self-administration studies). **Nardos Kebede & Jennifer Sedaille:** Writing – review & editing, Data curation (Vapor self-administration studies). **Tore Eid:** Writing – review & editing, Data curation (Blood nicotine and cotinine measurements). **Hanno C. Erythropel & Julie B. Zimmerman:** Writing – review & editing, Data curation (E-liquid chemical analysis). **Nii A. Addy:** Writing – review & editing Supervision, Funding acquisition, Conceptualization, Resources.

## Ethics approval

Animal experiments were conducted in accordance with The Yale University Institutional Animal Care and Use Committee (IACUC) guideline (IACUC Protocol no. #2022-11366)

## The list of nonstandard abbreviations

VSA: vapor self-administration
PG: propylene glycol
G: glycerol

